# PERK/ATF3-dependent induction of GDE4 modulates intracellular lysophospholipid-PPARα/γ signaling

**DOI:** 10.64898/2026.08.27.747495

**Authors:** Keisuke Kitakaze, Risa Misumi, Saya Nagai, Hanif Ali, Yu Ukai, Daisuke Takamine, Naoya Takehara, Yume Iiboshi, Rikuto Miyoshi, Yuko Ito, Yoshihiro Sunada, Yasuhiro Takenouchi, Kazuhito Tsuboi, Tamotsu Tanaka, Yasuo Okamoto

**Affiliations:** Department of Pharmacology, Kawasaki Medical School, Kurashiki, Okayama 701-0192, Japan; Department of Medical Technology, Kawasaki University of Medical Welfare, Kurashiki, Okayama 701-0193, Japan; Graduate School of Technology, Industrial and Social Sciences, Tokushima University, Tokushima 770-8513, Japan; Department of Anesthesiology and Intensive Care Medicine, Kawasaki Medical School, Kurashiki, Okayama 701-0192, Japan; Department of Neurosurgery 1, Kawasaki Medical School, Kurashiki, Okayama 701-0192, Japan

## Abstract

Lysophosphatidic acid (LPA) is widely recognized as an extracellular lipid mediator; however, the functional significance of intracellularly produced LPA remains poorly understood. Here, we investigated the regulatory mechanism and functional role of a LPA-producing lysophospholipase D GDE4, also known as GDPD1, in prostate cancer cells. GDE4 expression is induced under ER stress conditions in a PERK-dependent manner and requires the transcription factor ATF3. Disruption of GDE4 expression resulted in altered intracellular levels of LPA and LPA precursor lysophosphatidylethanolamine, accompanied by reduced cell proliferation. RNA sequencing and subsequent validation identified a set of genes downregulated in GDE4-depleted cells. Pharmacological inhibition experiments indicated that peroxisome proliferator-activated receptor α and γ (PPARα and PPARγ) signaling pathways contribute to the regulation of these GDE4-dependent genes. Collectively, our findings suggest that GDE4-dependent lipid remodeling is associated with PPARα/γ-mediated transcriptional regulation under ER stress conditions. These results provide a potential framework for understanding the link between intracellular lipid metabolism and stress-responsive gene regulation.

**Graphical abstract:** 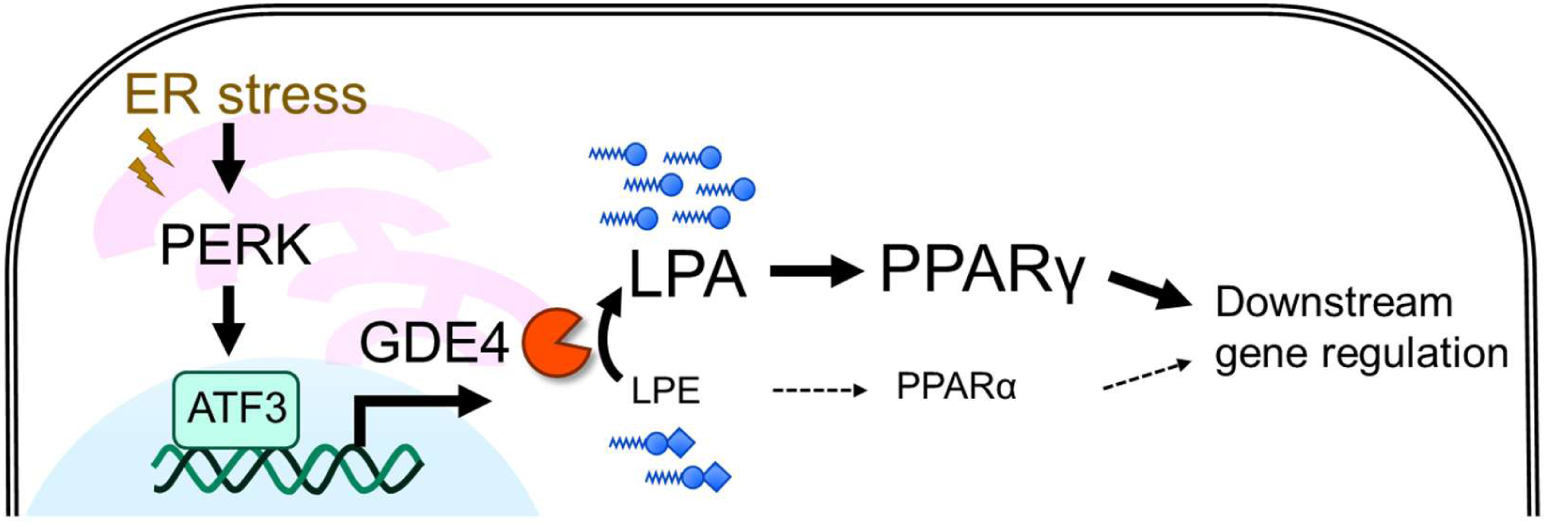

## Introduction

Lysophosphatidic acid (LPA) is a bioactive lipid present in eukaryotic tissues and plasma, mainly produced by the autotaxin (ATX), a lysophospholipase D (lysoPLD)-type enzyme in plasma [1]. Extracellularly produced LPA from lysophosphatidylcholine (LPC) functions as a lipid mediator via cell-surface LPA receptors, thereby regulating various biological processes including cell proliferation, migration, adhesion, platelet aggregation, inflammatory responses, and cancer invasion and metastasis [2]. On the other hand, glycerophosphodiesterase GDE4 and GDE7 —also known as GDPD1 and GDPD3, respectively— are localized to the endoplasmic reticulum (ER) membrane [3, 4] and act as intracellular lysoPLDs with active sites in the lumen [5, 6], producing LPA from lysophospholipids including LPC and lysophosphatidylethanolamine (LPE) [3, 4, 7, 8]. However, their functional roles and signaling pathways remain poorly understood. The lysoPLD enzyme GDE7 is involved in PPARγ-mediated signaling [6, 9]; however, it remains unclear whether the LPA produced by GDE4 functions as an intracellular signaling molecule. Furthermore, neither its target receptors nor downstream effectors have been identified. Recently, the expression of GDE4 and GDE7 was reportedly increased under stress conditions such as hypoxia, resulting in increased LPA production [10]. However, the underlying regulatory mechanisms and functional consequences have not been sufficiently elucidated. Therefore, the functional significance of GDE4 for intracellular LPA metabolism and signaling remains unclear.

The unfolded protein response (UPR) is a critical adaptive response mechanism that counteracts disturbances in protein homeostasis under ER stress conditions. The UPR consists of three major signaling pathways mediated by PERK, IRE1α, and ATF6 [11]. These pathways regulate cell survival and fate decisions through translational suppression and transcriptional regulation and play important roles in a variety of pathological conditions, including cancer [12]. Although UPR regulates lipid metabolism [13], its relationship with intracellular LPA production and GDE4 expression remains unclear.

This study aimed at clarifying the regulatory mechanism of GDE4 expression under ER stress conditions and examining whether GDE4-derived LPA contributes to intracellular signaling pathways associated with gene expression.

## Results

### ER stress induces GDE4 expression in a PERK pathway-dependent manner

First, we examined the regulatory mechanisms of GDE4 expression. Human prostate cancer LNCaP cells, which highly express endogenous GDE4 [14], were treated with ER stress inducers such as thapsigargin, tunicamycin, and dithiothreitol. Under nearly all treatment conditions, the transcript levels of GDE4 (*GDPD1*) and GDE7 (*GDPD3*) increased significantly, concomitant with the induction of CHOP (*DDIT3*) and BiP (*HSPA5*), the established UPR markers [11] (Fig. 1A).

**Figure 1.**
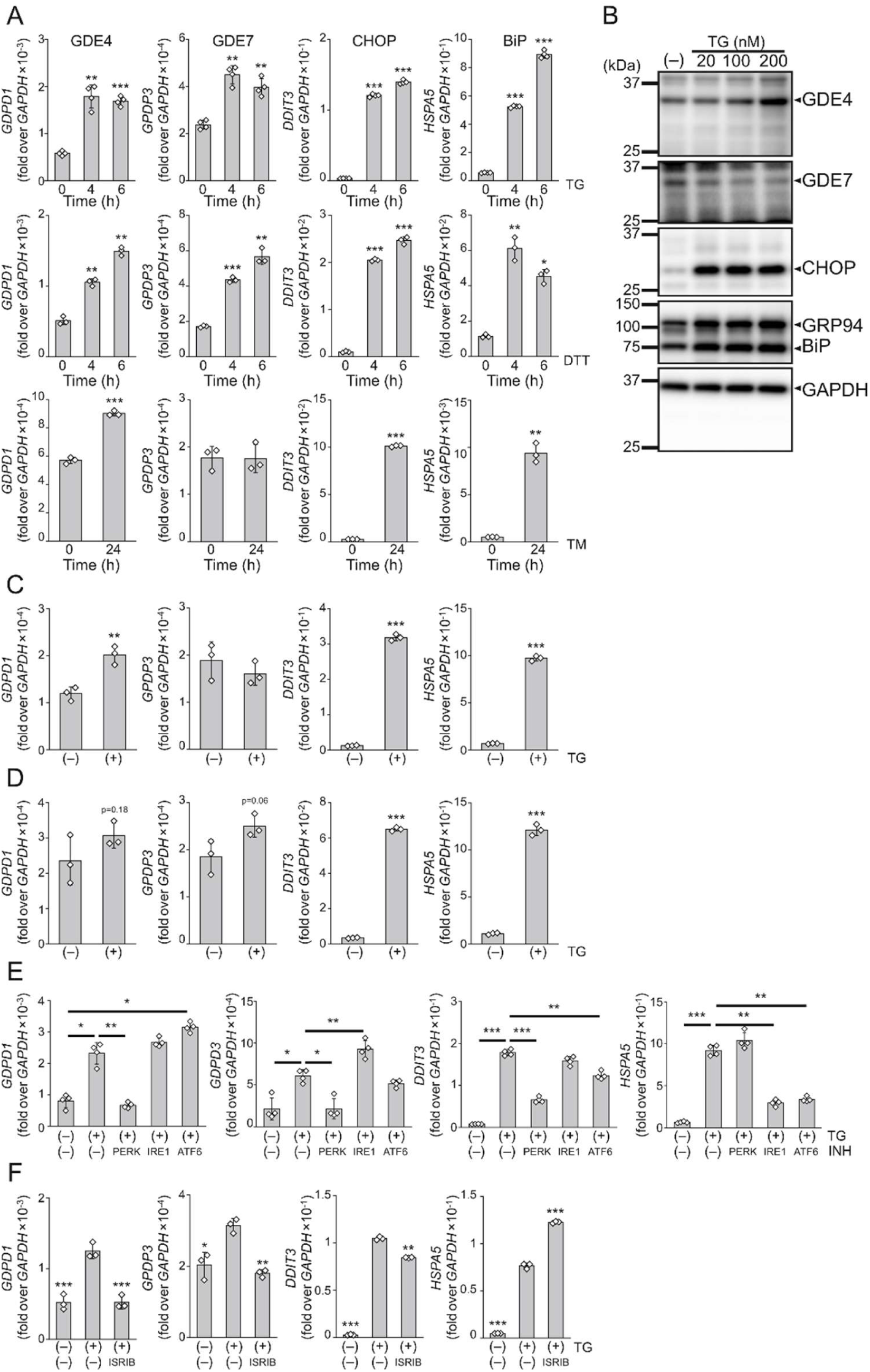
ER stress induces GDE4 expression in a PERK-dependent manner. (A) LNCaP cells were treated with ER stress inducers (200 nM thapsigargin [TG] or 10 mM dithiothreitol [DTT] for 0–6 h, or 2 μg/mL tunicamycin [TM] for 24 h). mRNA levels were analyzed by qPCR (n = 3 or 4). (B) Protein levels of GDE4 in LNCaP cells treated with 0–200 nM TG for 24 h were assessed by immunoblot analysis. (C, D) Induction of GDE4 mRNA expression by ER stress in prostate cancer cell lines PC3 (C) and DU145 (D) following treatment with 200 nM TG for 6 h (n = 3). (E) LNCaP cells were treated with 200 nM TG for 6 h in the presence of GSK2606414 (PERK inhibitor), APY29 (IRE1α inhibitor), or PF-429242 (ATF6 inhibitor), and GDE4 mRNA expression was analyzed by qPCR (n = 4). (F) Effect of ISRIB on GDE4 expression in LNCaP cells treated with 200 nM TG for 6 h (n = 3). Data are presented as mean ± SD. Statistical significance was determined by one-way ANOVA followed by Dunnett’s test. *p <0.05, **p <0.01, ***p <0.001.

Consistently, ER stress by thapsigargin led to an increase in GDE4 protein levels (Fig. 1B). Similar ER stress-mediated upregulation of GDE4 was observed in the other prostate cancer cell lines PC3 and DU145 (Fig. 1C, D). Next, to identify which UPR pathway is responsible for GDE4 induction, cells were treated with selective inhibitors of the PERK, IRE1α, and ATF6 pathways. Notably, ER stress-induced GDE4 upregulation was significantly suppressed only by the PERK inhibitor (Fig. 1E). Consistent with this finding, treatment with ISRIB [15], another pharmacological inhibitor of PERK pathway, also suppressed GDE4 induction. These results indicate that ER stress-induced GDE4 expression is dependent on the PERK pathway.

### Transcription factor ATF3 is required for induction of GDE4 expression

To elucidate the mechanism of PERK-dependent induction of GDE4 expression, we utilized ChIP-Atlas (https://chip-atlas.org/) to screen the transcription factors that could potentially bind to the 5′ untranslated region (5′UTR) of the human GDE4 gene. We identified ATF3 [16], a transcription factor of PERK pathway, as a candidate regulator of GDE4 expression (Fig. 2A). Similar to the altered expression of GDE4, the increase in ATF3 expression triggered by ER stress inducer was suppressed by PERK inhibitor and ISRIB treatment (Fig. 2B). To investigate the functional involvement of ATF3, we genome-edited the *ATF3* gene in LNCaP cells using the CRISPR-Cas9 system, and established a ATF3 knockout (KO) clone with heterozygous 2-bp and 5-bp deletion (Fig. 2C). In ATF3-KO cells, loss of ATF3 protein resulted in increased expression of *TMPRSS2* and *KLK3*, which are known androgen receptor (AR) target genes that are negatively regulated by ATF3 through suppression of AR signaling [17], confirming the functional disruption of ATF3 (Fig. 2D, E). Under ER stress condition, ATF3-KO cells retained robust induction of the UPR markers CHOP and BiP. However, the ER stress-induced GDE4 expression was almost abolished (Fig. 2E). Furthermore, we established the LNCaP cell lines in which the ATF3-binding site in the 5′UTR region of GDE4 was partially deleted (ΔBS1 and ΔBS2) (Fig. 2F). In these ΔBS cells, the expression levels of CHOP, BiP, and ATF3 were comparable to those observed in the wild-type (WT) cells, whereas GDE4 expression was significantly reduced both under basal and ER stress conditions (Fig. 2G). These results indicated that ATF3 binding to the 5′UTR of the GDE4 gene is required for both basal and ER stress-induced GDE4 expression.

**Figure 2.**
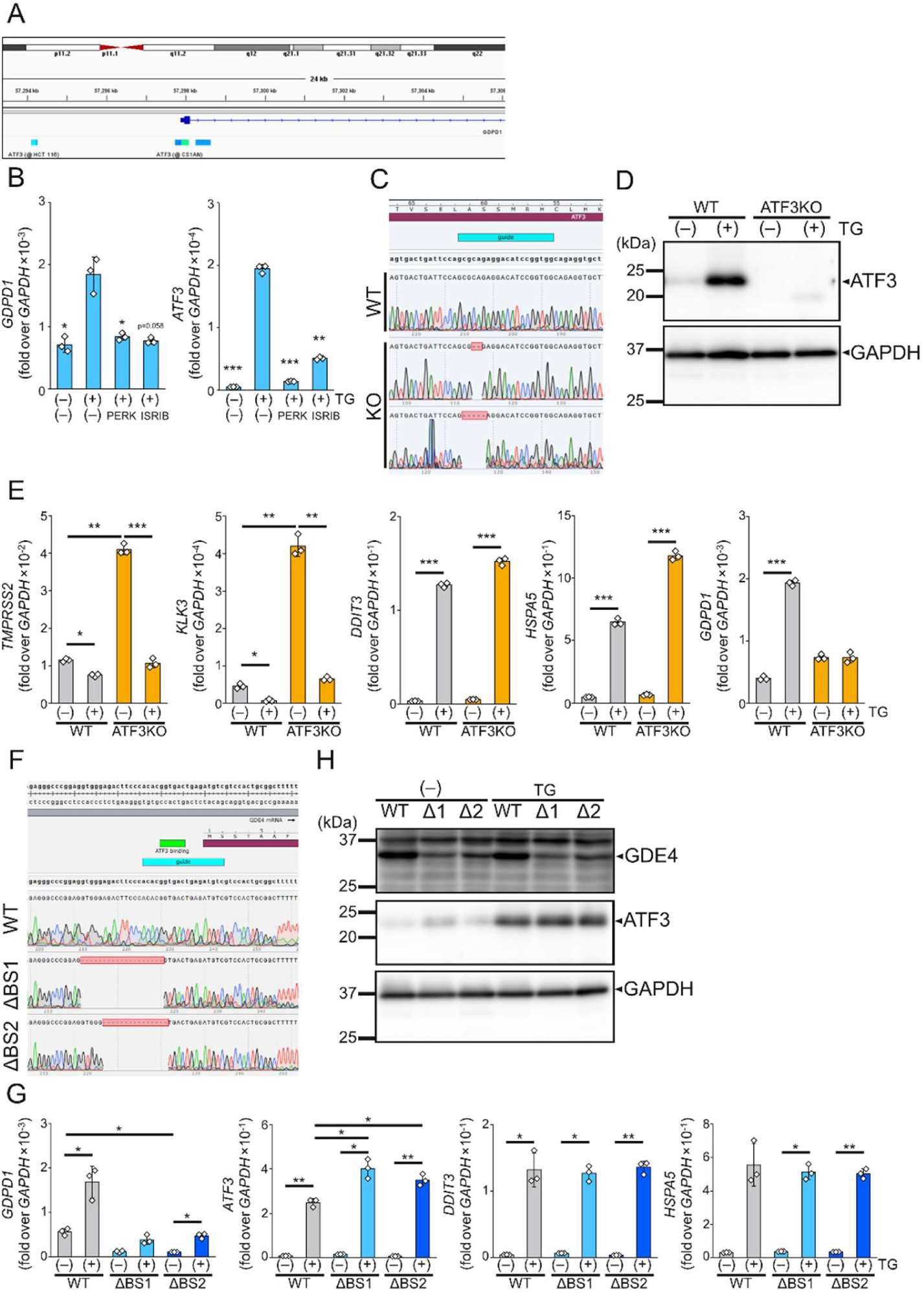
ATF3 is required for PERK-dependent induction of GDE4 expression. (A) Transcription factor binding profiles at the 5′ untranslated region (5′UTR) of the human GDE4 gene (*GDPD1*). (B) LNCaP cells were treated with 200 nM TG for 6 h in the presence or absence of PERK inhibitor or ISRIB, and mRNA expressions were analyzed by qPCR (n = 3). (C) Sequencing analysis of genome editing of the ATF3 gene in LNCaP cells, showing heterozygous 2-bp and 5-bp deletions in the ATF3-KO clone. (D) Immunoblot analysis of ATF3 protein in wild-type (WT) and ATF3-KO LNCaP cells treated with 200 nM TG for 24 h. (E) mRNA expressions in WT and ATF3-KO LNCaP cells treated with 200 nM TG for 6 h (n = 3). (F) Sequencing analysis of CRISPR-Cas9-mediated deletion of the ATF3-binding site in the 5′UTR of GDE4 gene, showing homozygous deletions of 19 bp (ΔBS1) and 15 bp (ΔBS2). (G) mRNA expressions in WT, ΔBS1, and ΔBS2 LNCaP cells under basal and ER stress conditions (200 nM TG, 6 h) (n = 3). (H) Immunoblot analysis of indicated proteins in WT, ΔBS1 (Δ1), and ΔBS2 (Δ2) LNCaP cells treated with 200 nM TG for 24 h. Data are presented as mean ± SD. Statistical significance was determined by one-way ANOVA followed by Dunnett’s or Tukey’s test. *p <0.05, **p <0.01, ***p <0.001.

### RNA-seq analysis identified candidate GDE4-regulated genes

We examined the phenotypes of disrupting GDE4 expression using ΔBS1/ΔBS2 cells, in which ATF3-dependent GDE4 induction is impaired, as well as previously established GDE4-KO cells [14]. GDE4-KO cells exhibited reduced proliferative capacity compared with WT cells (Fig. 3A), whereas these cells showed no significant changes in cell migration (Fig. S1), suggesting that GDE4 contributes to cellular signaling pathways involved in cell growth. To identify downstream factors modulated by GDE4, we performed RNA sequencing (RNA-seq) analysis. Hierarchical clustering revealed distinct gene expression profiles among WT, GDE4-KO, ΔBS1, and ΔBS2 cells (Fig. 3B). Differential expression analysis identified 6,267 significantly altered genes in WT vs. GDE4-KO, 3,988 in WT vs. ΔBS1, and 7,483 in WT vs. ΔBS2 (false discovery rate (FDR) <0.05) (Fig. 3C). Gene ontology (GO) enrichment analysis revealed significant changes in biological processes related to cytoplasmic translation, amino acid metabolism processes, chemical response, cell adhesion, and cell migration (Fig. 3D). Consistently, reactome pathway enrichment analysis showed significant enrichment of pathways related to translation, cell cycle, and electron transport chain (Fig. 3E). Among the differentially expressed genes, we focused on 17 genes (*CCDC85A*, *CNTNAP4*, *DHRS3*, *DNASE2B*, *DSC3*, *EPB41L3*, *GLP1R*, *H3-7*, *HS3ST3B1*, *IGHE*, *LOC100129434*, *MAGEA4*, *OSBPL7*, *PRDX3P1*, *STING1*, *TMSB4X*, and *ZNF608*) whose expression levels were consistently reduced to less than 0.5-folds in GDE4-KO, ΔBS1, and ΔBS2 cells compared to those in WT cells (Fig. 3F & Supplementary Table 1).

**Figure 3.**
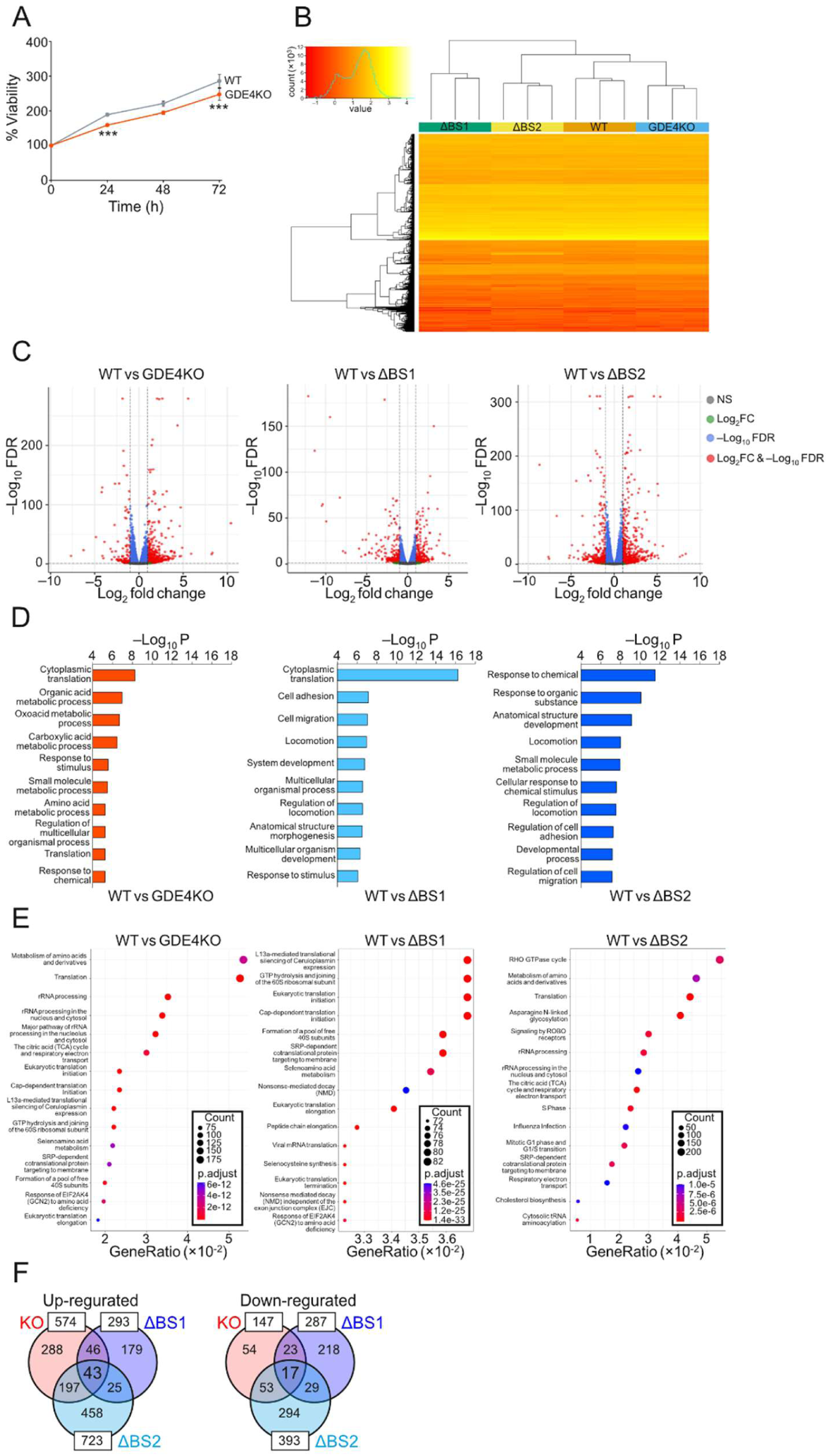
Identification of candidate GDE4-regulated genes by RNA-seq analysis. (A) Cell proliferation measured using the CellTiter-Glo assay. WT and GDE4 knockout (KO) LNCaP cells were assessed over 72 h in RPMI 1640 containing 1% charcoal-stripped FBS. Statistical significance was determined by unpaired *t*-test (***p <0.001). (B) Hierarchical clustering of RNA-seq data showing distinct gene expression profiles among WT, KO, ΔBS1, and ΔBS2 cells. (C) Volcano plots of differential gene expression in WT versus KO, WT versus ΔBS1, and WT versus ΔBS2 cells. NS indicates genes with no statistical significance; Log_2_FC represents genes with ≥ 2-fold change in expression; −log_10_FDR indicates genes with statistical significance (false discovery rate <0.05). (D) Gene Ontology (GO) enrichment analysis of differentially expressed genes in WT vs. KO, WT vs. ΔBS1, and WT vs. ΔBS2 cells. The top 10 enriched biological processes are shown. (E) Reactome pathway enrichment analysis of differentially expressed genes. The circle size represents gene count, and color indicates adjusted p-value (p.adjust). (F) Venn diagram showing the overlap of downregulated genes (≤ 0.5-fold vs. WT) among KO, ΔBS1, and ΔBS2 cells, highlighting 17 commonly downregulated genes. For RNA-seq analyses (B–F), cells were cultured in RPMI 1640 containing 1% charcoal-stripped FBS (n = 3).

### Validation and definition of GDE4-regulated hit genes

Next, we performed quantitative PCR (qPCR) to validate the commonly down-regulated genes (Fig. 4A), excluding the pseudogene *PRDX3P1* and the non-coding RNA *LOC100129434*. First, *TMSB4X* and *ZNF608* exhibited statistically significant and reproducible decreases, confirming the robustness of the RNA-seq findings. These genes were considered high-confidence downstream targets of GDE4. Second, *MAGEA4* showed statistically significant differences; however, these results were not consistently reproduced, suggesting potential context-dependent regulation. Third, *CNTNAP4*, *DSC3*, and *H3-7* displayed a consistent trend toward decreased expression, although statistical significance was not consistently achieved across comparisons. These genes may represent putative targets with reasonably high confidence. Finally, the other genes did not show a consistent decrease in expression by qPCR despite being identified in the RNA-seq dataset. These discrepancies may reflect differences in sensitivity between RNA-seq and qPCR as well as biological variability due to low gene expression levels. We also confirmed that the protein levels of DSC3, MAGEA4, and ZNF608 were reduced in GDE4-KO, ΔBS1, and ΔBS2 cells compared to those in WT cells (Fig. 4B). Six genes, namely *CNTNAP4*, *DSC3*, *H3-7*, *MAGEA4*, *TMSB4X*, and *ZNF608*, were then defined as hit genes and subjected to further analysis.

**Figure 4.**
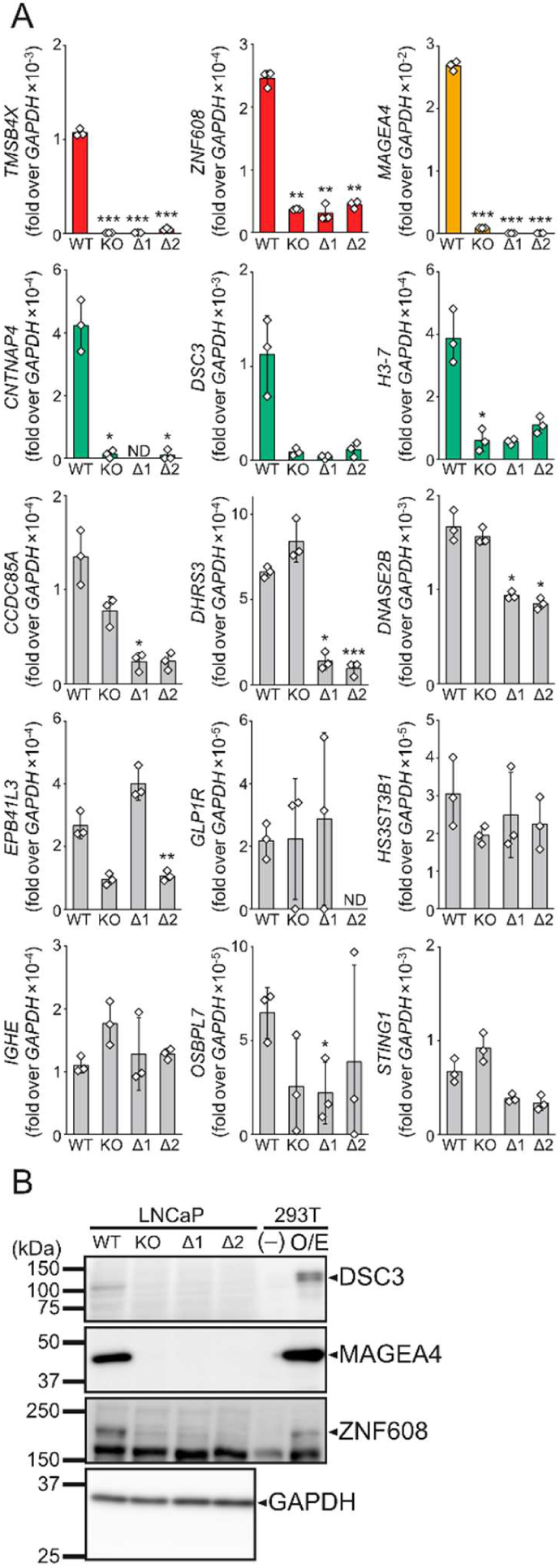
Validation and definition of GDE4-regulated hit genes (A) qPCR validation of downregulated genes identified by RNA-seq analysis in WT, GDE4 knockout (KO), ΔBS1 (Δ1), and ΔBS2 (Δ2) LNCaP cells. Genes are grouped based on reproducibility and statistical significance. Red, green, yellow, and gray bars indicate high-confidence, moderate-confidence, poorly reproducible, and unchanged genes, respectively. Data are presented as mean ± SD (n = 3). Statistical significance was determined by one-way ANOVA followed by Dunnett’s test. MAGEA4 showed statistically significant differences; however, these changes were not consistently reproduced across experiments. *p <0.05, **p <0.01, ***p <0.001. ND indicates not detected. (B) Immunoblot analysis of representative proteins, including DSC3, MAGEA4, and ZNF608, in WT, KO, Δ1, and Δ2 cells. O/E indicates overexpression. All experiments were performed in RPMI 1640 containing 1% charcoal-stripped FBS.

### GDE4 contributes to LPA production

Free fatty acids are known ligands for the nuclear receptors PPARα and PPARγ [18]. LPA acts not only through LPA receptors but also as a ligand for PPARγ [19], while LPC has been reported to act on PPARα [20]. Based on these observations, we investigated whether GDE4 contributes to intracellular LPA production. To this end, we quantified the levels of LPC and LPE, the substrates of GDE4, as well as the product LPA in WT and GDE4-KO cells using mass spectrometry. The LPC levels were not markedly altered, whereas the LPE levels were elevated in GDE4-KO cells. The intracellular LPA levels were significantly reduced in GDE4-KO cells compared to those in WT cells (Fig. 5A). These results are consistent with the previously reported substrate specificity of mouse GDE4 [3], and suggest that GDE4 contributes to intracellular LPA production primarily through the degradation of LPE rather than LPC. Next, we examined whether GDE4 deficiency led to changes in the expression of LPA receptors and PPARs. Consistent with previous studies [21], LNCaP cells expressed types 2 and 3 LPA receptors as well as PPARα and PPARγ (Fig. 5B), and their expression levels showed limited changes under the tested conditions.

**Figure 5.**
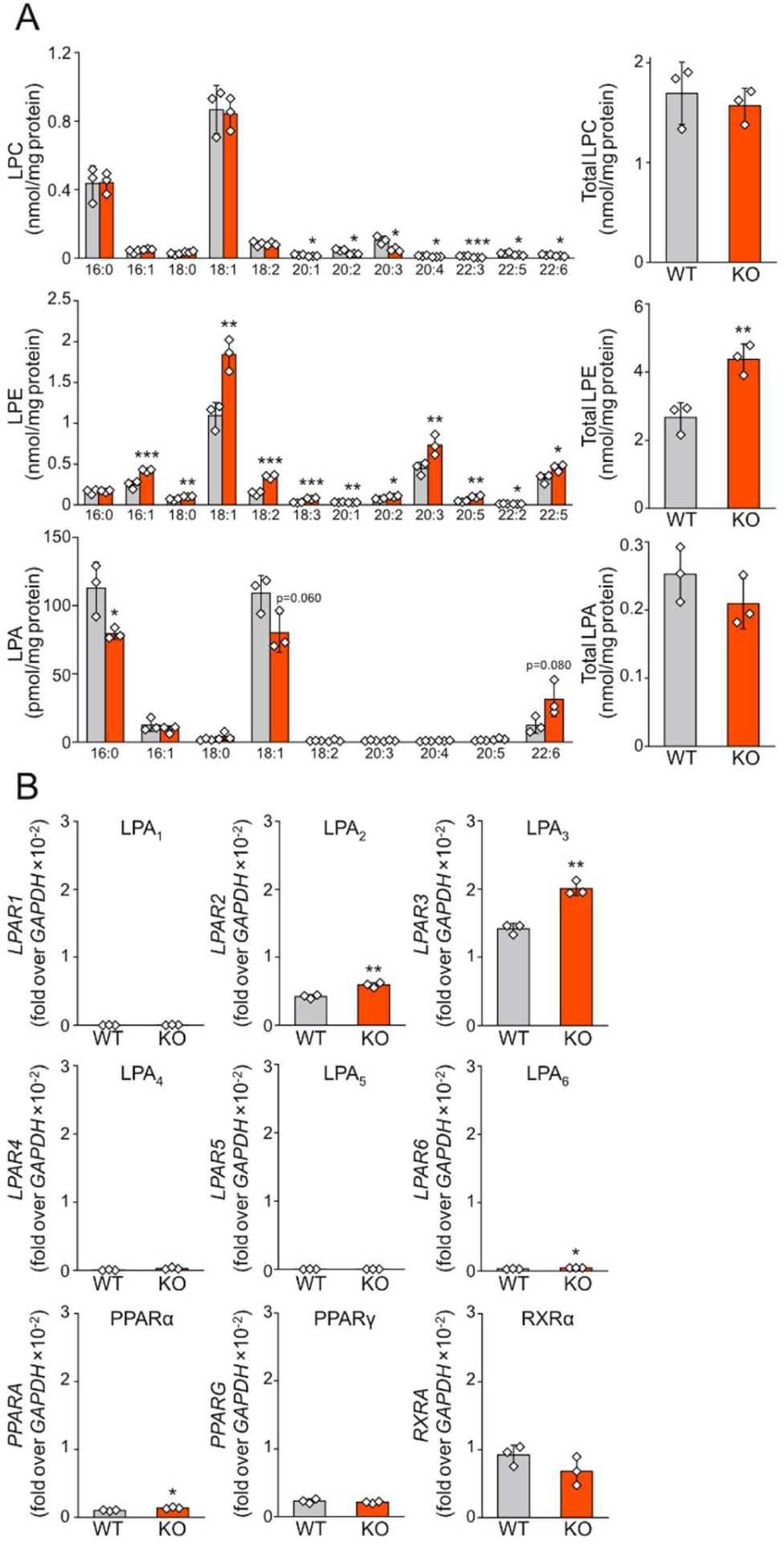
GDE4 contributes to intracellular LPA production. (A) Mass spectrometry-based quantification of LPC, LPE, and LPA species in WT and GDE4 knockout (KO) LNCaP cells. Individual molecular species and total levels are shown. Data are presented as mean ± SD (n = 3). Statistical significance was determined by unpaired t-test. *p <0.05, **p <0.01, ***p <0.001. (B) mRNA expression levels of LPA receptors (LPAR1–6) and nuclear receptors PPARα, PPARγ, and RXRα in LNCaP cells were analyzed by qPCR (n = 3). Statistical significance was determined by unpaired *t*-test. *p <0.05, **p <0.01.

### GDE4-dependent regulation of downstream gene expression is associated with PPARα/γ signaling

To identify receptors involved in GDE4-dependent downstream gene regulation, WT LNCaP cells were treated with antagonists targeting individual receptors. The expression levels of hit genes were then analyzed. Treatment with the LPAR2 antagonist H2L5186303 [22] and LPAR1/3 antagonists Ki16425 [23] had minimal effects on hit gene expression (Fig. 6A, B). In contrast, treatment with the PPARα antagonist GW6471 [24] resulted in increased expression of *DSC3* and *MAGEA4*, while significantly reducing the expression of *ZNF608* (Fig. 6C). Similarly, treatment with the PPARγ antagonist SR16832 [25] led to marked downregulation of *CNTNAP4*, *H3-7*, *TMSB4X*, and *ZNF608* (Fig. 6D). To determine whether extracellular LPA could regulate these genes, we treated WT and GDE4-KO cells with exogenous LPA. However, LPA treatment caused only minor changes in the expression of the hit genes (Fig. S2). In contrast, treatment of GDE4-KO cells with XY-4—an LPA derivative with selective PPARγ agonist activity [19]—resulted in a significant recovery in the expression of *TMSB4X* and *H3-7* (Fig. 6E). These findings support the model in which intracellular LPA derived from GDE4, rather than extracellular LPA signaling, regulates the expression of downstream genes via PPARγ signaling. Finally, we examined whether ER stress-induced regulation of hit gene expression depends on GDE4. Upon ER stress induction by thapsigargin, the expression levels of the UPR-related genes *DDIT3*, *HSPA5*, and *ATF3* increased significantly in both WT and GDE4-KO cells. In contrast, hit genes were markedly altered in WT cells, whereas distinct responses were observed in GDE4-KO cells (Fig. 6F). These results suggest that under ER stress conditions, GDE4-dependent alterations in LPA and LPE levels are associated with the regulation of the hit gene cluster, potentially involving PPARα-and PPARγ-mediated signaling pathways.

**Figure 6.**
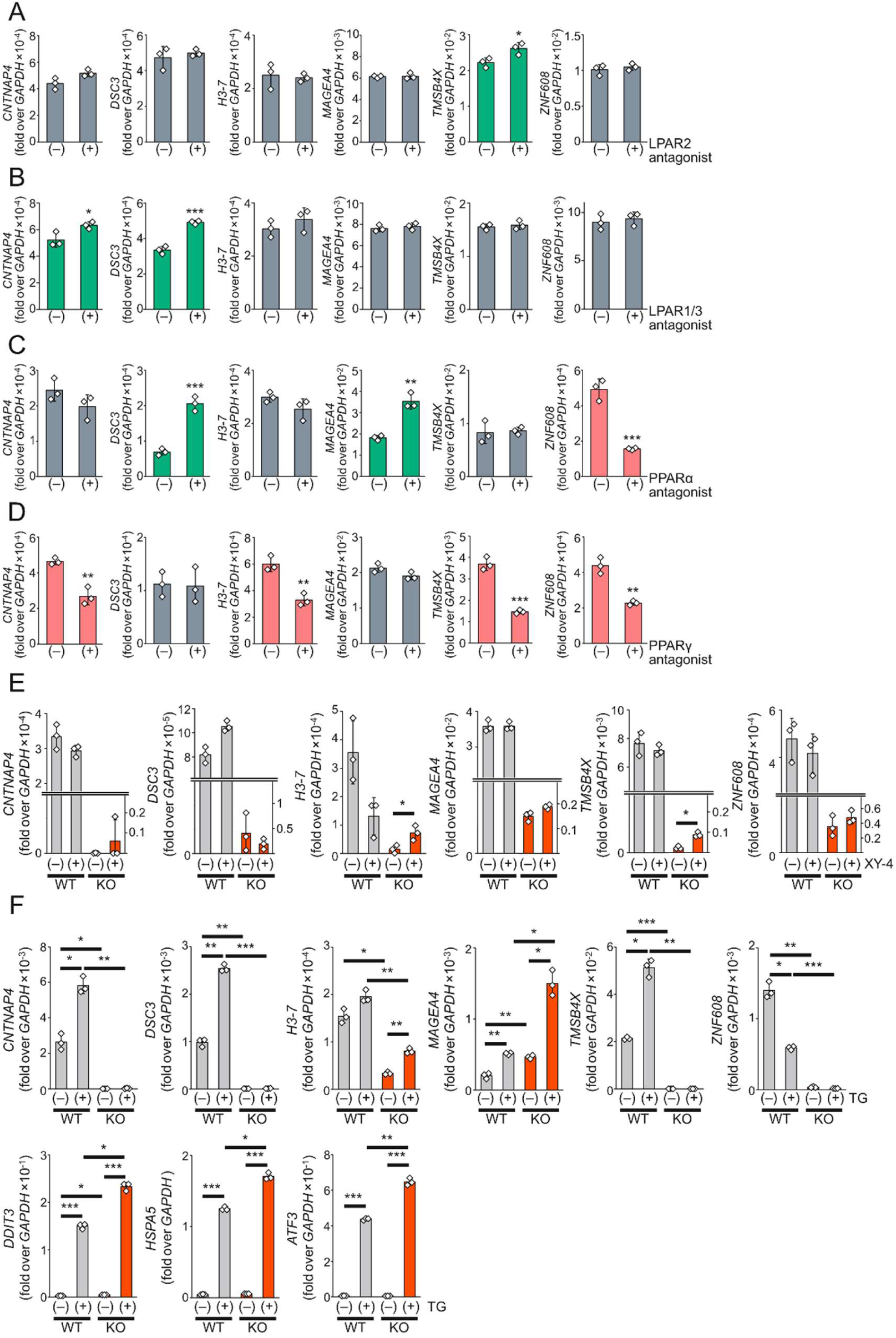
The GDE4-PPARα/γ pathway regulates downstream gene expression. (A–D) WT LNCaP cells were treated with antagonists targeting LPA or nuclear receptors for 24 h, and the expression of hit genes was analyzed by qPCR. Cells were treated with H2L5186303 (LPAR2 antagonist) (A), Ki16425 (LPAR1/3 antagonist) (B), GW6471 (PPARα antagonist) (C), or SR16832 (PPARγ antagonist) (D). Statistical significance was determined by unpaired *t*-test. (E, F) WT and GDE4 knockout (KO) LNCaP cells were treated with a PPARγ agonist XY-4 or ER stress inducer thapsigargin (TG) (F) for 24 h, and the expression levels of hit genes were analyzed by qPCR. Statistical significance was determined by one-way ANOVA followed by Tukey’s test. Data are presented as mean ± SD (n = 3). *p <0.05, **p <0.01, ***p <0.001. All experiments were performed in RPMI 1640 containing 1% charcoal-stripped FBS.

## Discussion

In this study, we demonstrated that GDE4 is induced by ER stress in a PERK pathway-dependent manner and that the transcription factor ATF3 is essential for this transcriptional regulation. We further showed that GDE4 contributes to changes in intracellular LPE and LPA levels, and that these alterations are associated with the expression of downstream gene sets via PPARα and PPARγ. These findings suggest that LPA, which has been predominantly understood as an extracellular signaling molecule, may also function as an intracellular signaling molecule.

Although intracellular signaling elicited by extracellular LPA through cell-surface LPA receptors has been extensively studied, the physiological significance of intracellularly generated LPA remains poorly understood. Several enzymes other than ATX and GDE4 are capable of producing LPA, including GDE7 and glycerol-3-phosphate acyltransferase (GPAT1). In addition to producing LPA, GDE7 preferentially produces cyclic phosphatidic acid (cPA) [6], a reported endogenous inhibitor of PPARγ [26], and has been implicated in PPARγ-related signaling. GPAT1 also generates intracellular LPA and activates PPARγ [27], but its LPA production occurs via a distinct biosynthetic route from glycerol-3-phosphate. These observations suggest that intracellular lysophospholipid metabolism may influence nuclear receptor signaling through multiple lipid mediators and biosynthetic pathways.

In the present study, the reduction of intracellular LPA levels caused by GDE4 deficiency was accompanied by the loss of downstream gene expression. These findings suggest that at least a subset of intracellular LPA may function as a signaling molecule involved in transcriptional regulation. An important implication of this study is that stress-induced GDE4 expression may shift the signaling balance from the LPE-PPARα pathway toward the LPA-PPARγ pathway, thereby contributing to the regulation of specific gene sets. In GDE4-KO cells, intracellular LPE levels were increased while LPA levels were reduced, whereas in WT cells, ER stress induced GDE4-dependent changes in the expression of hit genes. These results suggest that GDE4 may function not merely as an LPA-producing enzyme but also as a switch that modulates the balance of nuclear receptor-mediated transcriptional regulation. Consistent with the reduced proliferative capacity observed in GDE4-KO and ΔBS cells, several of the identified downstream genes, including *TMSB4X* [28, 29], *CNTNAP4* [30], and *DSC3* [31], are involved in cell adhesion and cell proliferation. Further studies will be required to clarify the phenotypic consequences mediated by these target genes. Also, whether GDE4-derived LPA can act on other LPA receptors that are not expressed in LNCaP cells remains to be investigated.

The lipid metabolic regulation has been frequently associated with the IRE1α-XBP1s pathway [32]. In contrast, PERK pathway is well known as a central mediator of translational suppression and stress-adaptive gene expression [33]. Our findings reveal that the PERK-ATF3 axis also regulates lipid metabolism through the expression of an LPA-producing enzyme. ATF3 is induced not only by ER stress but also by various forms of cellular stress as part of the integrated stress response [34]; thus, it is conceivable that the GDE4-LPA pathway is activated under a broad range of stress conditions. Accordingly, the GDE4-LPA pathway may function as a component of a general cellular stress adaptation mechanism rather than being restricted to ER stress. From this perspective, the GDE4-LPA axis may function not merely as a downstream effector of the PERK pathway, but as a metabolic arm of stress signaling that links stress sensing to the remodeling of intracellular lipid composition and nuclear receptor-mediated transcriptional programs. Such mechanism could allow cells to coordinately adjust both protein homeostasis and lipid signaling in response to stress. Further studies will be required to clarify how changes in lipid balance driven by GDE4 contribute to the maintenance of ER homeostasis.

Although GDE4 has been implicated in retinal degeneration [35], its involvement in diseases remains poorly understood. Recently, GDE4 was identified as a metabolism-related hub gene associated with biochemical recurrence following radical prostatectomy, suggesting a potential role for GDE4 in prostate cancer progression [36]. In contrast, GDE7 regulates cancer recurrence via cancer stem cells [37] and promotes epithelial-mesenchymal transition and prostate adenocarcinoma progression through the LPA/LPAR1/AKT pathway [38]. These observations raise the possibility that GDE4 may also play important roles in various pathological conditions. These findings and our results suggest that abnormal intracellular LPA production may exert broad pathological effects beyond the traditionally recognized role of extracellular LPA signaling. In addition, our data support a model in which GDE4-dependent modulation of intracellular LPA levels is associated with PPAR-dependent transcriptional programs that influence cellular phenotypes such as proliferation and adhesion.

Several limitations of the present study should be considered. First, although our findings support an association between GDE4-dependent LPA production and PPARα/γ-mediated transcriptional regulation, direct evidence that GDE4-derived LPA activates these nuclear receptors is still lacking. The partial restoration of hit gene expression by the PPARγ agonist XY-4 supports the involvement of PPARγ signaling, but does not establish that GDE4-derived LPA is the endogenous ligand responsible for this effect. Direct assessment of PPARα/γ activation and genetic approaches targeting these pathways will be required to establish this causal relationship. Second, the mechanistic analyses were primarily performed in LNCaP cells. Although ER stress-induced GDE4 expression was also observed in PC3 and DU145 cells, whether the GDE4-PPARα/γ axis is conserved across different prostate cancer subtypes or other cell types remains unknown. In addition, the present study is primarily based on *in vitro* cell models, and thus the generalizability of these findings to *in vivo* systems remains to be established.

In summary, this study proposes a lipid signaling model in which ER stress-induced GDE4 is associated with intracellular LPE/LPA balance and controls PPARα/γ-dependent gene expression. Our findings provide insights into the interplay between lipid metabolism and transcriptional regulation during cellular stress adaptation and establish a conceptual framework for understanding the potential roles of GDE4 in diverse pathological conditions, such as cancer, fibrosis, and degenerative diseases.

## Methods

### Cell line

Human embryonic kidney HEK293T cells were maintained in DMEM supplemented with 10% (v/v) FBS (Biofluids) and nonessential amino acids. Human prostate cancer cell lines LNCaP (RIKEN BRC, RCB2144), PC-3 (RIKEN BRC, RCB2145), and DU145 (RIKEN BRC, RCB2143) were maintained in RPMI 1640 medium supplemented with 10% (v/v) FBS and 2 mmol/L L-alanyl-L-glutamine. Human GDE4 knockout LNCaP cells were established as previously described [14]. All cell lines were cultured at 37 °C in a humidified atmosphere containing 5% CO_2_, and were confirmed to be mycoplasma-free using the MycoBlue Mycoplasma Detector (Vazyme Biotech).

### Plasmid construction and cell transfection

pcDNA3.1(+) plasmid vectors harboring the cDNA of human DSC3, MAGEA4, and ZNF608 with a C-terminal FLAG tag was constructed by Genscript. HEK293T cells were cultured to 80% confluence in a poly-L-lysine-coated 6-well plate. The cells were then transfected with 4.5 μg of the plasmid using 5.6 µL Lipofectamine 2000 (Thermo Fisher Scientific). Control cells were also prepared using an insert-free vector. At 48 h post-transfection, the cells were harvested using the lysis buffer.

### Genome editing

Specific guide RNA sequences were selected for the human *ATF3* and 5′UTR of GDE4 gene using CRISPOR [39] and cloned into the LentiGuide-Puro vector (gift from Dr. Feng Zhang, Addgene plasmid #52963). The resultant LentiGuide-*ATF3*-Puro and LentiGuide-5′UTR-Puro plasmids, as well as lenti-Cas9-Blast (gift from Dr. Feng Zhang, Addgene plasmid #52962), were used to prepare the lentivirus as previously described [40]. Briefly, HEK293T cells were co-transfected with pMD2.G and psPAX2 (gifts from Dr. Didier Trono, Addgene plasmids #12259 and #12260, respectively) together with each plasmid using Lipofectamine 2000, and lentivirus-containing conditioned media were collected. LNCaP cells were transduced using the conditioned media containing lenti-Cas9-Blast lentivirus in the presence of 10 μg/mL hexadimethrine bromide and subjected to selection using 10 μg/mL blasticidin S. Cas9-expressing LNCaP cells were subsequently transduced using the conditioned media containing LentiGuide-*ATF3*-Puro and LentiGuide-5′UTR-Puro lentivirus sequentially and positive clones were selected using 2 μg/mL puromycin. The clonal cell lines were isolated using the limiting dilution method. Genomic DNA from each clone was amplified and sequenced using an ABI 3130 DNA sequencer or SeqStudio Genetic Analyzer (Thermo Fisher Scientific-Applied Biosystems) for verification. The used primer pairs are listed in Supplementary Table 2.

### ER stress induction and chemical treatment

For ER stress induction, cells were seeded in 6-well plates and treated with 200 nM thapsigargin, 2 μg/mL tunicamycin, or 10 mM dithiothreitol. Where indicated, cells were co-treated with thapsigargin and either 1 μM PERK inhibitor I (GSK2606414, Merck), 2.5 μM APY-29, 2 μM PF-429242, or 200 nM ISRIB (Cayman). An equivalent volume of DMSO was added to control cells. After incubation at 37 °C for 6 or 24 h, total RNA was extracted.

For analysis of downstream signaling, cells were cultured in RPMI 1640 medium containing 1% charcoal-stripped FBS (Sigma, F6765) for 48 h to minimize the influence of ATX and LPA present in FBS. For antagonist treatment, at 24 h after seeding, the medium was replaced with fresh medium containing one of the following antagonists: 20 μM H2L5186303 (LPAR2 antagonist, Cayman), 20 μM Ki16425 (LPAR1/3 antagonist, ChemScene), 50 μM GW6471 (PPARα antagonist, Cayman), 50 μM GW9662 (PPARγ antagonist, Cayman), 2 μM SR16832 (PPARγ antagonist, Cayman), 50 μM LPA (Echelon Biosciences), or 10 μM XY-4 (PPARγ agonist, Echelon Biosciences). Cells were further cultured for 24 h, and total RNA was then extracted.

### Immunoblot analysis

Cells were lysed in lysis buffer (50 mM Tris–HCl (pH 7.5), 150 mM NaCl, 2 mM EDTA, 1% Nonidet P-40, and 0.1% sodium dodecyl sulfate [SDS]) with Halt Protease and Phosphatase Inhibitor Cocktail (Thermo Fisher Scientific) and were centrifuged at 12,000 *g* for 15 min at 4 °C. The supernatants were then used as cell lysates. Protein concentration was determined with the Protein Assay BCA Kit (Nacalai Tesque) using BSA as a standard. The obtained samples were separated on SDS-polyacrylamide gel and electrotransferred onto polyvinylidene difluoride (PVDF) membranes (Immobilon-E, Merck Millipore). The PVDF membranes were blocked with phosphate-buffered saline (PBS) containing 0.1% Tween 20 (PBS-T) and 5% skim milk (Morinaga Milk Industry) for 1 h at 25 °C. They were then probed with anti-GDE4 (1:1500 dilution in 2% BSA/PBS-T, Proteintech, 27861-1-AP), anti-GDE7 (1:1000, Atlas Antibodies, HPA041148), anti-CHOP (1:1000, Proteintech, 15204-1-AP), anti-KDEL (1:5000, Medical & Biological Laboratories, M181-3MS), anti-ATF3 (1:1000, Cell Signaling Technology, 33593), anti-DSC3 (1:1000, PROGEN, 61093), anti-MAGE-A4 (1:1000, Cell Signaling Technology, 82491), anti-ZNF608 (1:1000, Atlas Antibodies, HPA005545), and anti-GAPDH (1:1000, Medical & Biological Laboratories, M171-3) for 18 h at 4 °C. After the membranes were washed three times with PBS-T, the bound antibodies were visualized using horseradish peroxidase-linked anti-rabbit or anti-mouse IgG secondary antibody (1:1000, Cell Signaling Technology, 7074S and 7076S) depending on the primary antibody. Protein expression signals were detected with Pierce ECL western blotting substrate (Thermo Fisher Scientific), and the chemiluminescent signals were quantified using an Amersham Imager 680 (Cytiva).

### Lipid analysis of cultured cells by liquid chromatography-tandem mass spectrometry (LC-MS/MS)

Cellular lipids were extracted using the Bligh and Dyer method [41] with internal standards. The following synthetic standards were added prior to extraction: C17:0 LPE (100 pmol), C17:0 LPA (100 pmol), and C17:0 LPC (100 pmol). The extracted lipids were dissolved in 100 μL of chloroform/methanol/water (1:2:0.2, v/v/v) and subjected to non-targeted lipidomic analysis. Chromatographic separation was performed using an Acquity UPLC BEH-C18 column (50 × 2.1 mm, 1.7 μm; Waters, Ireland) coupled to an Ultimate 3000 UHPLC system (Thermo Fisher Scientific). The flow rate was maintained at 0.3 mL/min, and the column temperature was set at 45 °C. Mobile phase A consisted of acetonitrile/methanol/water (1:1:3, v/v/v) containing 5 mM ammonium acetate and 10 nM EDTA, whereas mobile phase B consisted of isopropanol containing 5 mM ammonium acetate and 10 nM EDTA. The injection volume was 5 μL, and the autosampler temperature was maintained at 4 °C. The gradient program was as follows: 0–1 min, 0% B; 5 min, 40% B; 7.5–12 min, 64% B; 12.5 min, 82.5% B; 19 min, 85% B; 20 min, 95% B; and 20.1–25 min, 0% B. Mass spectrometric analysis was performed using a Q Exactive Orbitrap mass spectrometer (Thermo Fisher Scientific). Data were acquired using data-dependent acquisition (DDA), with MS1 collected at a resolving power of approximately 35,000 full width at half maximum (FWHM) and MS2 at approximately 20,000 FWHM. The mass range for both MS1 and MS2 scans was m/z 70–1250. Lipid species were identified and quantified using MS-DIAL metabolomics software [42]. Quantification was performed by calculating the ratio of endogenous lipid peak areas to those of the corresponding internal standards.

### RNA isolation and quantitative PCR (qPCR)

RNA isolation, cDNA synthesis, and qPCR were performed as previously described [43]. Briefly, RNA was isolated from the cells using TRIzol reagent (Thermo Fisher Scientific). The quality and concentration of the isolated RNA were estimated by measuring the absorbance at 260 and 280 nm on a Nanodrop One spectrophotometer (Thermo Fisher Scientific). cDNA was then synthesized with PrimeScript RT Master Mix (TaKaRa Bio). The relative gene expression was quantified by qPCR using a TB Green Premix Ex Taq II (TaKaRa Bio), which was performed using a StepOnePlus or QuantStudio5 Real-Time PCR System (Applied Biosystems) in accordance with the manufacturer’s protocols. The used primers are listed in Supplementary Table 2. An average threshold cycle (Ct) value was calculated and normalized to that of the housekeeping gene *GAPDH* to obtain the ΔCt value.

### Viability and migration assays

LNCaP cells were seeded on a 96-well plate and then incubated for 0–72 h. Cell viability was measured using CellTiter-Glo (Promega) with VarioskanFlash. Cell migration was evaluated using the CytoSelect 24-Well wound healing assay kit (Cell Biolabs) according to the manufacturer’s instructions.

### RNA-seq

Sequencing libraries were generated using NEBNext Ultra RNA Library Prep Kit for Illumina (NEB, Cat No. 7760) following the manufacturer’s protocol, and index codes were added to attribute sequences to each sample. The resulting libraries were purified (AMPure XP system), and library quality was assessed on the Agilent Bioanalyzer 2100 system. The purified libraries were sequenced on a Novaseq 6000 (Illumina) with a 150-base pair-end read setting. Sequence quality was checked using FastQC [44] and MultiQC [45] with --export option. Trimmomatic [46] was used to remove adapter sequences, and PRINSEQ-lite [47] was used to remove the poly A/T tail, short sequence length reads, and trim low-quality bases at the extremity of the reads. Reads were aligned to the Homo sapiens reference genome (hg38) using STAR [48]. Gene expression levels were calculated using the featureCount function from the Subread [49], and differentially expressed genes (DEGs) were determined using the R package edgeR [50]. In DEG analysis, gene expressions were normalized to counts per million (cpm), and genes expressed in less than 2 samples were removed. Expression levels after TMM normalization were used for DEG analysis. Hierarchical clustering was performed for genes expressing more than 1 cpm in at least 2 samples using R packages amap and gplots. For clustering, the Euclidean distance was calculated, and the complete linkage method was used. Gene Ontology (GO) enrichment analysis was performed using the R package topGO [51] with the “classic” method and Fisher’s Exact test. Reactome Pathway enrichment analysis was performed using the R package ReactomePA [52]. The database org.Hs.eg.db [53] was used for each enrichment analysis.

## Statistics and Reproducibility

All data are expressed as the mean ± SD. Statistical analyses were performed using Prism (version 8.42; GraphPad Software). The sample sizes from one-time experiment, reproducibility, and number of biological replicates used are indicated in the Figure legends. Unpaired, two-tailed Student’s *t*-test was used to compare between two groups, while ANOVA followed by Dunnett’s or Tukey’s post-hoc test was used to compare between three or more groups. P-values <0.05 were considered as statistically significant.

## Data availability

All data provided in the article and Supplementary files are available from the corresponding author upon reasonable request.

## Acknowledgments

The authors would like to thank Rie Sugimoto for technical assistance and also acknowledge technical support from Central Research Institute, Kawasaki Medical School. The authors thank Editage (www.editage.com) for English language editing. This work was supported by the Japan Society for the Promotion of Science KAKENHI (grant numbers JP23K14810, JP25K14827 to K.K.); ONO Medical Research Foundation to K.K.; the Uehara Memorial Foundation (Grant Number 202310014 to K.K.); Teraoka Scholarship Foundation to K.K.; Wesco Scientific Promotion Foundation to K.K.; the KAWASAKI Foundation of Medical Science and Medical Welfare to K.K.; EA Pharma Co., Ltd. (grant number AS2022A000003641 to K.K. and Y.O.); Santen Pharmaceutical Co., Ltd. (grant number SPCS20220415001 to K.K. and Y.O.); Teijin Nakashima Medical Co., Ltd. (to K.K. and Y.O.); Asahi Kasei Pharma Corporation (grant number APJS20230401001 and APJS20250408006 to K.K. and Y.O.); CSL Behring K.K. (grant number AS2022A000009767 to K.K. and Y.O.); Taiho Pharmaceutical Co., Ltd. (AS2023A000002034 to K.K. and Y.O.); Zimmer Biomet G.K. (ZBCS20250408004 to K.K. and Y.O.), and Research Project Grants from Kawasaki Medical School (grant numbers R04Y-001, R05B-067, R07B-070, and R08I055 to K.K.). The funders had no role in study design, data collection, decision to publish, or preparation of the manuscript.

## Author contributions

Conceptualization, K.K.; Formal analysis, K.K.; Investigation, K.K., R.M., S.N., H.A., Y.U., D.T., N.Y., Y.Iiboshi, and R.M.; Data curation, K.K.; Writing - Original Draft, K.K. and Y.O.; Writing - Review & Editing, K.K., H.A., Y.Ito, Y.S., Y.T., K.T., and T.T.; Visualization, K.K.; Project administration; K.K.; Funding acquisition, K.K. and Y.O.; All authors approved the manuscript for publication.

## Declaration of interests

The authors declare the following competing interests: K.K. and Y.O. received grants from EA Pharma Co., Ltd., Santen Pharmaceutical Co., Ltd., Teijin Nakashima Medical Co., Ltd., Asahi Kasei Pharma Corporation, CSL Behring K.K., Taiho Pharmaceutical Co., Ltd., Zimmer Biomet G.K. All other authors declare no conflicts of interest.

